# Histone H3.3 Hira chaperone complex contributes to zygote formation in mice and humans

**DOI:** 10.1101/2020.06.18.159954

**Authors:** Rowena Smith, Sue Pickering, Anna Kopakaki, K Joo Thong, Richard A Anderson, Chih-Jen Lin

## Abstract

Elucidating the underlining mechanisms underpinning successful fertilisation is imperative in optimising IVF treatments, and may lead to a specific diagnosis and therefore potential treatment for some infertile couples. One of the critical steps involves paternal chromatin reprogramming, in which compacted sperm chromatin packed by protamines is removed by oocyte factors and new histones, including histone H3.3, are incorporated. This step is critical for the formation of the male pronucleus, without which the zygote contains only 1 pronucleus (1PN), in contrast to normally fertilised zygotes with two-pronuclei (2PN). 1PN zygotes are a frequently observed phenomenon in IVF treatments, therefore aberrant mechanism of action controlling paternal chromatin repackaging may be an important cause of abnormal fertilisation. Hira is the main H3.3 chaperone that governs this protamine-to-histone exchange. In this study, we investigated the maternal functions of two other molecules of the Hira complex, Cabin1 and Ubn1 in the mouse. Loss-of-function Cabin1 and Ubn1 mouse models were developed: their zygotes displayed abnormal 1PN zygote phenotypes, similar to the phenotype of Hira mutants. We then studied human 1PN zygotes, and found that the Hira complex was absent in 1PN zygotes which were lacking the male pronucleus. This result confirms that the role of the Hira complex in male pronucleus formation has coherence from mice to humans. Furthermore, rescue experiments showed that the abnormal 1PN phenotype derived from Hira mutants could be resolved by overexpression of Hira in the mouse oocytes. In summary, we have provided evidence of the role of Hira complex in regulating male pronucleus formation in both mice and humans, that both Cabin1 and Ubn1 components of the Hira complex are equally essential for male pronucleus formation, and that this can be rescued. We present a proof-of-concept experiment that could potentially lead to a personalised IVF therapy for oocyte defects.

## Introduction

Fertilisation of an oocyte by a sperm to produce a zygote and then an embryo underlies mammalian reproduction. As a treatment for infertility, IVF is well established, but although fertilisation remains at its centre, our understanding of this process is limited, preventing the possibility of specific diagnoses and thus treatments that may be relevant to some couples, and which may improve success rates overall. To achieve a successful fertilisation event, consequential steps including oocyte activation, chromatin reprogramming of the sperm (protamine-to-histone exchange and histone reassembly), and formation of the pronuclei are essential. These all rely on the quality of the oocytes, and this is dependent on oocyte stored factors (Gosden, 1996) (Swain and Pool, 2008). Therefore, it is imperative to elucidate the underlining regulatory mechanisms involved in these steps, in the search for potential new IVF treatments which can overcome defects found in poor quality oocytes. An intriguing clinical phenotype arising from abnormal fertilisation is the one-pronucleus (1PN) zygote. It is observed after overnight culture following both conventional IVF and ICSI and occurs in 3%-17% of fertilised oocytes, though mostly in the range of 4-8% (Azevedo et al., 2014; Si et al., 2019). However, the etiology is unknown.

The building blocks of chromatin are the core histones H2A, H2B, H3, and H4. In addition to these canonical histones, there are variants which share similar sequences with canonical histones and which carry out diverse functions. One example is histone variant H3.3, a version of H3 with wider regulatory roles, one of which, significantly, is that it allows H3.3 to bind to chromatin independently of the cell cycle (Maze et al., 2014). The incorporation of histones onto chromatins is tightly regulated by histone chaperones (Hammond et al., 2017), (Buschbeck and Hake, 2017). Hira complex, which comprises Hira, Cabin1, and Ubn1, is one of the H3.3 chaperones (Rai et al., 2011). Applying a mouse genetic model, we have previously reported that maternal Hira is essential for the completion of chromatin reprogramming of the sperm during fertilisation: mutant Hira oocytes formed abnormal 1PN zygotes that lacked a male pronucleus due to the failure of H3.3 incorporation (Lin et al., 2014). The maternal roles of the other molecules in the Hira complex, Cabin1 and Ubn1, remain unknown.

In this study, we explored the maternal role of the remaining candidates within the Hira complex, Ubn1 and Cabin1, in the mouse. We have found that they are essential in the paternal chromatin reprogramming event. Loss of function mouse models for either Ubn1 or Cabin1exhibited the same 1PN phenotypes as Hira mutants, and rescue experiments revealed that the abnormal 1PN phenotype of Hira mutants could be restored back to 2PN zygotes. Our examination of abnormal human 1PN zygotes showed that failure of Hira complex binding is implicated in their formation. This sheds light on the potential for rescuing maternal factors in humans as an approach to personalised Assisted Reproductive Technology (ART) medical therapy.

## Results

### Maternal Hira complex is involved in male chromatin deposition in mouse oocytes

Firstly, we examined whether the constituent molecules of the Hira complex were present in wild type mouse oocytes during oogenesis. Immunohistochemistry of mouse ovarian sections demonstrated that both Cabin1 and Ubn1 were present in oocytes and were particularly enriched in germinal vesicle GV nuclei (Fig. 1A). Wholemount immunofluorescence of GV stage oocytes also confirmed this observation. Therefore, both Cabin1 and Ubn1 are oocyte factors that accumulate in GV oocytes.

**Figure1.**
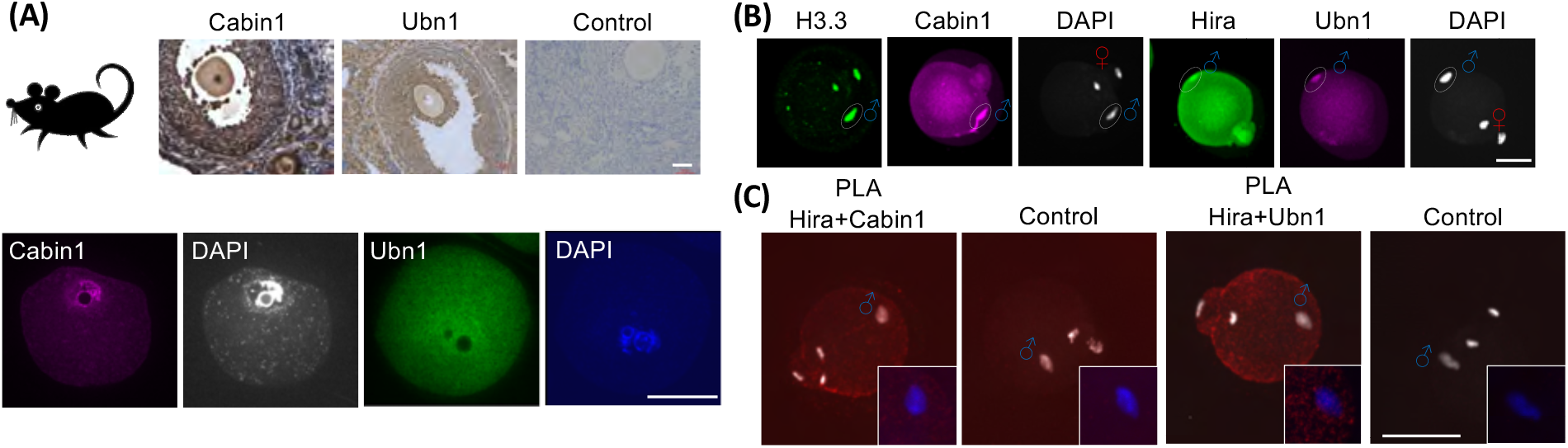
Maternal Hira complex incorporates into male chromatin after fertilisation. (A) Cabin1 and Ubn1 are maternal factors present in the mouse oocytes. Immunohistochemisty of murine ovarian sections (upper panel) and immunofluorescence of GV oocytes (lower panel) showed positive signals of Cabin1 and Ubn1 in the GV nuclei. Scale bar=50μm. (B) Cabin1 and Ubn1 bind to male chromatin after fertilisation. Immunofluorescence of H3.3 and Cabin1 (left panel) and Hira and Ubn1 (right panel) show positive signals in the decondensed male chromatin after fertilisation. Scale bar=50μm (C) Proximity ligation assay (PLA) confirmed the interactions of Cabin1with Hira (upper panel) and Ubn1with Hira (lower panel) in the male chromatin during sperm decondensation. Upper panel: Images of embryos stained with antibodies against Hira and Cabin1. Negative control: antibody of hCG and Cabin1; Lower panel: Images of embryos stained with antibodies against Hira and Ubn1. Negative control: antibody of hCG and Ubn1; Inset: Enlarged (1.7 fold) images of male chromatins. Original to 30cm and crop Scale bar=50μm.

We postulated that both Cabin1 and Ubn1 collaborate with Hira and are involved in the chromatin reprogramming processes during fertilisation. We collected and fixed newly fertilised IVF embryos (2-4 hours post fertilisation) in which female chromatin had completed the 2nd meiotic division and in doing so had segregated sister chromatids. Meanwhile, male chromatin had undergone DNA decondensation. Immunofluorescence showed that Cabin1 had deposited into chromatin and co-localised with H3.3. Ubn1 had also deposited into male chromatin and showed a similar pattern to Hira [Fig. 1B; (Lin et al., 2014)].

To provide further evidence of the interactions within Hira complex during male chromatin decondensation, we performed proximity ligation assay (PLA), a sensitive assay for visualisation of protein -protein interactions. We detected positive signals for Hira and Cabin1 distributed throughout the zygotes, and for Hira and Ubn1 in the enriched foci in the decondensed male chromatin, but not in the negative controls (Fig. 1C). Thus, we demonstrated that both Cabin1 and Ubn1 are maternal factors deposited in oocytes and incorporated into decondensed male chromatin after fertilisation and that they physically interact with Hira. We hypothesised that the molecules within the Hira complex could collaboratively play a role during paternal chromatin reprogramming.

### Hira complex is essential for male pronucleus formation during fertilisation in mice

We then generated three new Hira complex models, each one with oocyte loss-of-function in one component part of the Hira complex (Fig. S1A).

We generated a new Hira conditional knockout mouse line different from our previous report (Lin et al., 2014), (Lin et al., 2013). We used a Zp3-Cre mouse line to inactivate the Hira flox/flox alleles and so delete exon 6-7. We observed an infertile phenotype in the new Hira mutant mice identical to that seen in our previous model (data not shown). Zygotes retrieved from this new Hira mutant also displayed 1 PN phenotypes due to the failure of male pronucleus formation [Fig. 2A].

**Figure 2.**
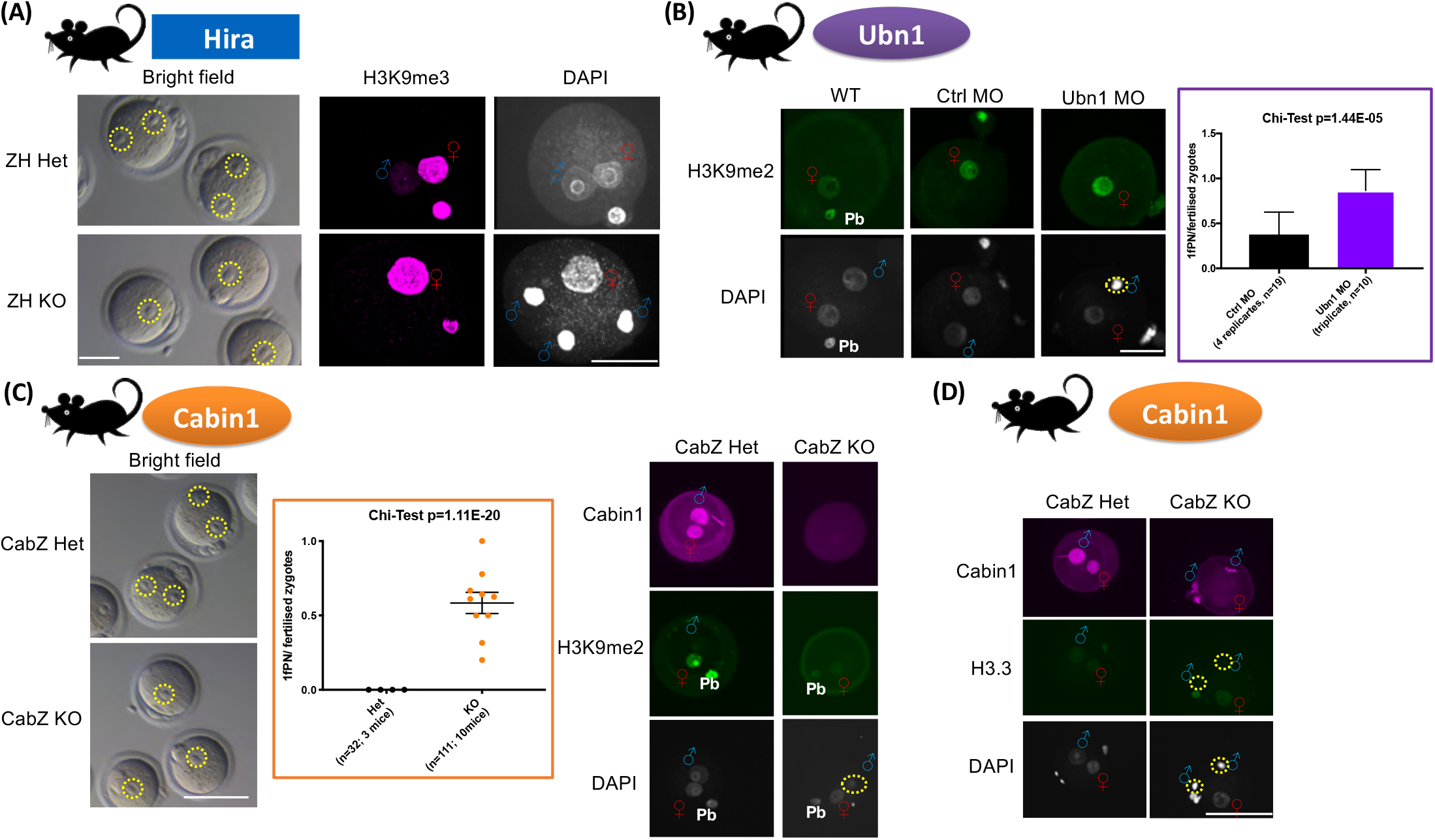
Loss-of-function maternal Hira complex in mice gave rise to abnormal one-pronucleus (1PN) zygotes due to failure to form male pronuclei. (A) Hira mutant oocytes formed abnormal 1PN zygotes after fertilisation. Brightfield images showed abnormal 1PN zygotes of Hira mutants (ZH KO), in contrast to normal two-pronuclei (2PN) zygotes in the controls (ZH Het; left panel). Immunofluorescence showed that male pronucleus formation is impaired in Hira mutant zygote (right panel). Female chromatin was labelled with histone H3K9me3. Scale bar=40μm. (B) Ubn1 knockdown oocytes formed 1PN zygotes after fertilisation. Male pronucleus formation is impaired in Ubn1 morpholino (MO) knocked down zygotes. Histone H3K9me2 stained for female chromatin (left panel). Abnormal 1PN formation rate was increased in the Ubn1 MO knockdown zygotes (right panel). Scale bar=40μm. (C) Cabin1 mutant oocytes formed abnormal 1PN zygotes after fertilisation. Brightfield images showed abnormal 1PN zygotes formed in the Cabin1 mutants (CabZ KO), in contrast to normal 2PN zygotes in the controls (CabZ Het; left panel). Abnormal 1PN formation rate was increased in the Cabin1 mutant zygotes (middle panel). Immunofluorescence showed that male pronucleus formation is impaired in the Cabin1 mutants. Female chromatin was labelled with histone H3K9me2 (right panel). Scale bar=80μm. (D)Histone H3.3 failed to incorporate into male chromatin in the Cabin1 mutant zygotes. Scale bar=80μm.

For Ubn1, we applied the morpholino antisense oligos knockdown approach to oocytes [(Lin et al., 2013; Lin et al., 2014); Fig. S1B]. After injection of morpholino oligos against Ubn1 (Ubn1 MO), the level of Ubn1 in oocytes was significantly reduced compared to control oligos (Ctrl MO) (Fig. S1C). Interestingly, fertilised Ubn1 MO matured oocytes revealed a high proportion (87%) of abnormal 1PN zygotes (Fig. 2B).

To investigate the maternal role of Cabin1, we again used the Zp3-Cre approach and crossed to Cabin1flox/flox mice to delete exon 6 (Fig. S1A). Cabin1 mutant mice (CabZ KO) also showed an infertile phenotype, and fertilised CabZ KO zygotes revealed the abnormal 1PN phenotype due to the failure of male pronucleus formation (Fig. 2C). Immunofluorescence on Cabin1 zygotes demonstrated that H3.3 was incapable of incorporating into male chromatin in CabZ KO oocytes. In contrast, H3.3 incorporated into pronuclei of Cabin1 heterozygous controls (CabZ Het) (Fig. 2D).

We also investigated whether the loss-of-function of Ubn1 and Cabin1 compromised the overall stability of the Hira complex. We performed immunofluorescence on Ubn1 knockdown oocytes and assessed the relative level of Hira and Cabin1, the two other molecules within the Hira complex. We found that the overall nuclear staining signals of Hira and Cabin1 were significantly decreased in Ubn1 MO oocytes compared to Ctrl MO oocytes (Fig. S2D).

Similarly, we conducted immunofluorescence on CabZ KO oocytes. Not only were overall nuclear staining signals of Ubn1 and Hira reduced in CabZ KO oocytes but so was the level of H3.3. We also performed the Triton-X pre-extraction protocol to remove the unbound chromatin associated signals in CabZ KO oocytes (Nashun et al., 2015), and noted that the level of H3.3 was comparably lowered in these oocytes (Fig. S1E).

The above results suggest that each molecule of Hira complex is essential for the stability of the complex in the oocyte, and loss of any one of them affects male pronucleus formation at fertilisation.

### Hira complex failed to incorporate into male chromatin of human abnormal 1PN zygotes

Based on the promising data collected from mice, we investigated whether the Hira complex is critical for male pronucleus formation in humans.

Human 1PN embryos were obtained after donation for research use, after examination of zygotes 18 hours post insemination (including both IVF and ICSI). During the recruitment period of 2018-2019, we approached a total of 218 couples and 144 couples gave consent. Overall 23 1PN zygotes were identified from 16 IVF cycles and 21 were fixed (1PN /fertilised embryo: 21/101=20.1%; Table 1 and Fig. 3A). Subsequent analysis showed that an addition 9 embryos originally identified in the clinic as being 1PN were at more advanced stages upon collection. Seven had progressed to nuclear envelope breakdown (NEBD) or the 2-cell stage, and 2 were parthenogenetically activated.

**Figure 3.**
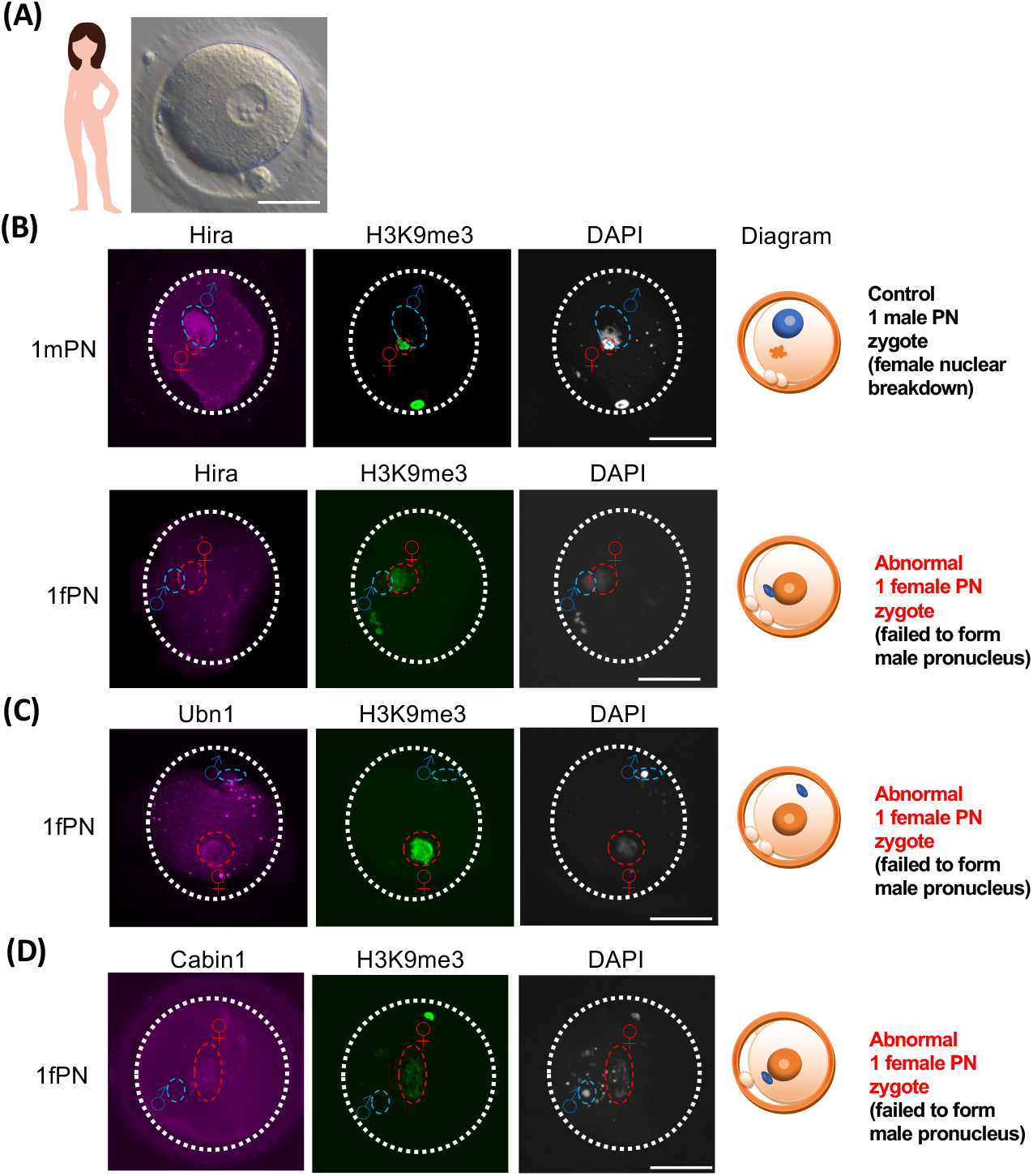
One-pronucleus (1PN) human zygotes revealed deficit of Hira complex incorporation into the male chromatin. (A) Representative image of a human 1PN zygote collected. (B) Hira failed to incorporate into the male chromatin of abnormalfemale 1PN zygote. Immunofluorescence of Hira and H3K9me3 of human 1PN zygotes. Hira incorporated into the male chromatin of a normally fertilised zygote (female chromatin progressed to nuclear breakdown, while male PN remained; upper panel), but failed to incorporate into the abnormal female 1PN zygote (failure of male pronucleus formation; lower panel). (C) Ubn1 failed to incorporate into the abnormal 1 female PN zygote. Immunofluorescence of Ubn1 and H3K9me3 of a human 1PN zygote. Ubn1 failed to incorporate into the male chromatin. (D) Cabin1 failed to incorporate into an abnormal 1 female PN zygote. Immunofluorescence of Cabin1 and H3K9me3 of a human 1PN zygote. Cabin1 failed to incorporate into the male chromatin of an abnormal 1female PN zygote. Histone H3K9me3 labels female chromatin; DNA counterstained by DAPI. Histone H3K9me2, a female chromatin specific marker. Scale bar=50μm

**Table 1.** Summary of the one-pronucleus zygote collection during recruitment period

| **Recruitment period** | **No. of patients approached** | **No. of patients consented** | **No. of patients showed 1PN zygotes** | **No. of 1PN zygotes collected** | **For experiments** |
| --- | --- | --- | --- | --- | --- |
| **2018** | **70** | **46** | **8** | **9** | **9** |
| **2019** | **148** | **98** | **8** | **14** | **12** |
| **Total** | **218** | **144** | **16** | **23** | **21\*** |

\*To note that not all zygotes were found to be 1PN on assessment in the laboratory, 9 were at other stages, specifically one-male pronucleus zygotes as Control x2; wrong stages x5, and parthenogenesis x2.

**Table 2.** Summary of immunofluorescence result of abnormal one-pronucleus zygote

| Stained antibodies | No. of one fPN zygotes stained with antibodies | Positive signal detected in male chromatin | Ratio |
| --- | --- | --- | --- |
| Hira | 4 | 0 | 0/4 |
| Ubn1 | 4 | 0 | 0/4 |
| Cabin1 | 4 | 2* | 2/4 |
| Total | 12 | 2 | 2/12 |

\*one with weak signal

We performed immunofluorescence on the fixed samples using H3K9me3, a female chromatin specific marker, in order to distinguish the origin of the parental chromatin. We discovered that there were two 1PN zygotes which were actually morphologically abnormal but cytologically normal diploid zygotes. The parental chromatins had developed asynchronously: the female chromatin had progressed to nuclear envelope breakdown stage, while the male chromatins had remained in the pronuclear stage [Fig. 3B; also observed in Egli et al 2011(Egli et al., 2011)]. In these 1 male PN zygotes, we observed that the Hira immunostaining signal was enriched (Fig. 3B). In contrast, in the abnormal 1female PN zygotes [female chromatin labelled by H3K9me3; (van de Werken et al., 2014)], the male chromatin had failed to form the male pronucleus, remained in the decondensation stage, and importantly lacked Hira staining. This is in agreement with the data collected from Hira mutant mice (Fig. 2A).

As in the loss-of-function Ubn1 fertilised 1PN phenotype in the mouse, we also found that Ubn1 was absent in the male chromatin of the 1PN human zygotes (Fig. 3C). We noted that in half of the 1PN zygotes we stained for Cabin1, signal was absent in the male chromatin (Fig. 3D). This was consistent with the abnormal 1PN formation rate in Cabin1 mutant mice (Fig. 2C). A summary of immunofluorescence results is shown in the Figure 3.

### Overexpression of Hira in mutant oocytes rescues male pronucleus formation in their zygotes

The results above show that maternal Hira complex is critical for male pronucleus formation and that loss-of-function leads to abnormal 1PN phenotypes in mice (Fig. 2). In these abnormal phenotypes, Hira complex fails to incorporate into male chromatin which is likely to contribute to the abnormal 1 female PN human zygote phenotype (Fig. 3). Both Hira and Cabin1 null mouse showed later developmental defects than maternal depletion ones (see discussion), thus, maternal deposition of Hira complex has a more profound role than previously understood during the oocyte-to-embryo transition, and this cannot be compensated by zygotic transcription rescue during the 2-cell stage.

Mitochondrial replacement therapy has been reported to be a substitution procedure for the rescue of oocyte inherited mitochondrial disease (Greenfield et al., 2017). We envisaged that abnormal 1PN zygote phenotypes caused by the maternal defect of Hira could potentially be rescued by overexpression in oocytes. To test this, we firstly adapted the same micromanipulation platform as we applied to knockdown of Ubn1 in oocytes (Fig. 4A). Instead of morpholino oligos (Fig. S1B), we injected in vitro transcribed RNA into Hira mutant (ZH KO) oocytes. The GFP-tagged protein was stably expressed, and we were able to increase Hira to a level comparable with that of Hira heterozygous (ZH Het) oocytes (Fig. S2).

**Figure 4.**
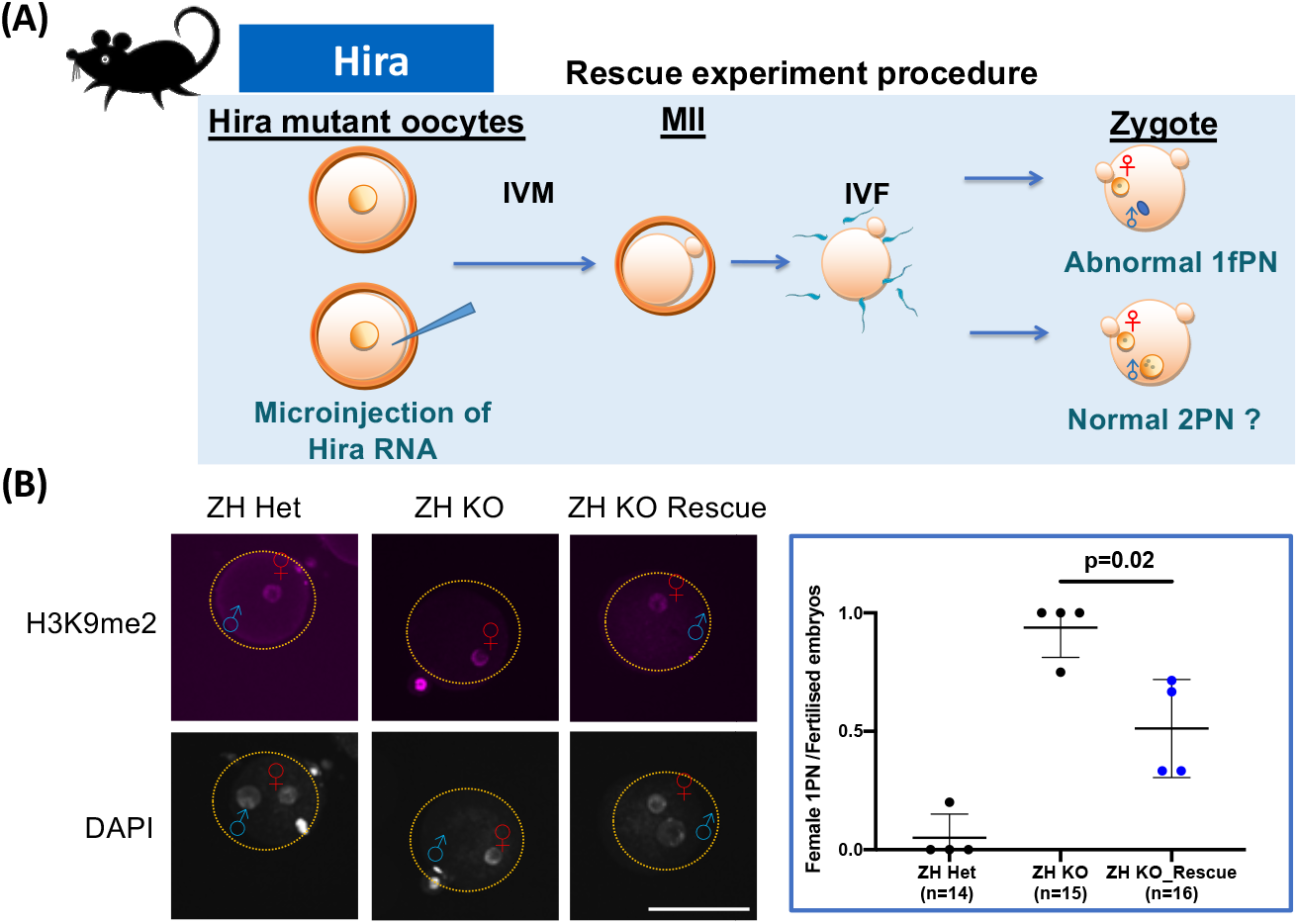
Overexpression of Hira rescued the male pronucleus formation in the Hira mutants. (A) Experimental procedure for rescuing maternal Hira mutants. Hira RNA was introduced into the GV oocytes, following maturation and fertilisation in vitro. Pronuclear formation was examined by immunofluorescence. (B) Over-expression of Hira RNA in the Hira mutant oocytes rescued the male pronucleus formation after fertilisation. Immunofluorescence of H3K9me2 showed that zygote with 2 pronuclei were formed after rescue (left panel). Summary of the rescue experiments (right panel). Scale bar=80μm.

We then introduced Hira RNA into Hira mutant oocytes. Following maturation and fertilisation we determined formation of the male pronucleus as an indicator of a successful rescue event by immunofluorescence (Fig. 4B). ZH KO fertilised embryos showed a high proportion of abnormal 1 female PN zygotes compared to ZH Het zygotes (86.7% versus 7%, p=2.7E-17). However, in ZH KO oocytes RNA injection partially rescued the proportion of abnormal 1 female PN zygotes from 86.7% to 56% (p=0.02; Fig. 4B). This result clearly supports our assertion that overexpression of Hira in null oocytes can support zygote formation, thus providing further support for the proof-of-principle approach to rescuing poor quality oocytes (here, null Hira) in order to support oocyte-to-embryo transition.

## Discussion

We have demonstrated, that in mice, Cabin1 and Ubn1 are maternal factors which are incorporated into decondensed male chromatin after fertilisation, and physically interact with Hira. We showed that molecules within the Hira complex, Cabin1 [a mammalian specific H3.3 chaperone (Orsi et al., 2013)] and Ubn1, acting interdependently, are also critical for male pronucleus formation in mouse. The roles of H3.3 and Hira have been reported in bovine pre-implantation stages (Zhang et al., 2016; Zhang et al., 2018), but relevant study, particularly focusing on maternal roles, has been lacking. Improved understanding will be beneficial for the improvement of domestic animal production. Importantly, our investigation of abnormal human 1PN zygotes showed that Hira complex was absent in the male chromatin, thus linking this unique phenotype with a coherent mechanism of zygote formation in mice and humans. Using our mouse model, we showed that we could ameliorate the deleterious effects of the loss of Hira complex, thus providing a possible approach for rescuing poor quality human oocytes for ART.

Results of molecular cytogenetics of human 1PN zygotes have been reported. These results show that the possible underlying causes include synchronous PN formation, fusion of maternal and paternal PN, parthenogenetic activation, or premature breakdown of PN (Azevedo et al., 2014). Among our limited but carefully-phenotyped 1PN samples (n=21) we observed that the majority showed failure of male pronucleus formation (Fig. 3), indicating an abnormal fertilisation event. Other zygotes, initially identified as having 1PN, were subsequently shown to have asynchronous PN formation (n=2; as the IF controls), parthenogenetic activation (n=2), or incorrect staging (progressed to nuclear breakdown and 2 cells; n=5) (Table 1). Our results support the statement that, in general, 1PN zygotes should be considered to have undergone abnormal fertilisation and to be less competent of progressing to a normal embryo (Destouni et al., 2018),(Yao et al., 2016),(Araki et al., 2018).

Originally, null mutants demonstrated the requirement for Hira around the time of gastrulation (Roberts et al., 2002) but our previous studies discovered that maternal Hira is essential for zygote development (Lin et al., 2014), (Nashun et al., 2015). Similarly, Cabin1 null mice showed embryo lethality around embryonic day 12.5 due to organogenesis defects (Esau et al., 2001). Interestingly, our oocytic deleted Cabin1 mutants revealed preimplantation defects (not shown). Based on the results of our Hira maternal mouse models, which revealed a great loss of developmental propensity (Lin et al., 2014), we envisaged that the lower developmental potential of human 1PN embryos might be due to defects of embryonic genome activation, arising at the 4-8 cell stages in humans. It would be valuable to measure the transcriptome of abnormal 1PN zygotes to establish a profile of the downstream effectors as a “good oocyte” signature. Also, it will be valuable to monitor the subsequent pregnant outcomes from patients who donated the 1PN zygotes and the reoccurrence of 1PN zygotes after new cycles.

Next we mined the human exon sequence dataset, GenomeAD Browser (https://gnomad.broadinstitute.org), to examine whether there is a genetic linkage relating to the loss-of-function of Hira complex. We noted that the loss-of-function observed/expected upper bound fraction (LOEUF) parameters of Hira (0.02-0.14) and Ubn1 (0.05-0.21) are very low [Cabin1 (0.36-0.59)], indicating low tolerance to inactivation of the genes. In other words, both Hira and Ubn1 likely tend to result in loss-of-function mutations. The interdependence of Hira complex components shown in Fig. S1D-E, Nashun et al (Nashun et al., 2015) and the previous data in human cells (Rai et al., 2011) indicates that the normal function of all three components of Hira complex are required for its adequate function. Similarly, the abnormal human 1PN zygotes we collected were possibly inherited from loss-of-function Hira complex oocytes. This means that abnormal 1PN zygotes might harbour divergent transcriptomic and chromatin profiles. Therefore, it will be of interest to collect human 1PN zygotes and perform RNA-sequencing or chromatin-associated assays [ATAC-seq (Wu et al., 2016) and Hi-C (Flyamer et al., 2017) or scNMT-seq (Clark et al., 2018)] to gain insight into human paternal genome reprogramming events.

Overexpression of Hira mRNA in mutant murine oocytes partially rescued the abnormal 1PN phenotype by significantly restoring mutant oocyte fertilisation to form normal 2 PN zygotes (Fig. 4). This not only demonstrated that maternally stored Hira is critical for male pronucleus formation but also provided a proof-of-principle experiment as the foundation of an approach for rescuing defects caused by impaired maternal factors. Analogous to mitochondria replacement therapy [MRT; (Greenfield et al., 2017)] we envisage that personalised medicine therapies could be applied in IVF fields in the near future. Our next steps will be to test the outcome of overexpression of other candidates such as Cabin1 or Ubn1 mutant mouse oocytes respectively, and possibly to develop nuclear transfer types of approach, such as GV transfer or spindle transfer(Costa-Borges et al., 2020), for subsequent advancement to an alternative MRT, 3 parents IVF, and other novel personalised medicine treatments. In the near future, a characterised maternal defect oocyte could potentially be rescued using a design platform for patients who have previously produced1PN zygotes after ART.

## Materials and methods

### Animals

Mouse experiments were approved by the University of Edinburgh’s Animal Welfare and Ethical Review Board (AWERB) and carried out under the authority of a UK Home Office Project Licence. The mouse lines used in this study carried conditional floxed alleles for Cabin1 and for Hira on C57BL/6 backgrounds (both were provided from Prof Peter Adams previously of the Beatson Institute, UK, now at Sanford Burnham Prebys, USA). Floxed Cabin1 (Cabin1 fl/fl) and floxed Hira (Hira fl/fl) mouse lines were crossed with a Zp3-Cre mouse line [(de Vries et al., 2000); provided from Prof Petra Hajkova, Imperial College of London] that expresses Cre recombinase in the female germline to generate heterozygous and homozygous mutant oocytes.

### Manipulation of Oocytes and embryos

Female mice were superovulated by administration of 7.5 I.U. pregnant mare serum gonadotrophin (PMSG from Prospec Protein Services, USA) or 80μl ultra-superovulation reagent (CARD HyperOva Cosmo Bio, Japan; (Hasegawa et al., 2016)) and 48 hours later fully-grown oocytes were isolated. For *in vitro* maturation (IVM), oocytes were cultured in M16 medium (Sigma) at 37°C, 6%CO2 and 5%O2 for 18 to 24 hours.

For zygote collection, 48 hours after superovulation, 7.5 I.U. human chorionic gonadotrophin (hCG, Chorulon® from Intervet) was injected and the mice mated with C57BL/6 males. The following day zygotes were collected from the oviducts and cumulus cells were removed after treatment with 300 units/mg hyaluronidase (Millipore).

The microinjection procedure was the same as previously described (Chen et al., 2013), (Lin et al., 2014). The micromanipulation platform was equipped with a microinjector (FemtoJet 4i, Eppendorf), an inverted microscope (Leica, DMi8), and micromanipulators (Narishige). For Ubn1 knockdown experiments, oocytes were microinjected with 2mM of either control or Ubn1 antisense morpholino oligos (Gene Tools).

For the Hira rescue experiments, Hira-GFP (Addgene 59781) and GFP (gift from M. Anger, IAPG, CAS) were used as templates for *in vitro* synthesis of RNA using mMessage mMachine® (Ambion, Life Technologies) followed by polyadenylation using a Poly(A) tailing kit (Applied Biosystems). Transcripts were purified by phenol:chloroform extraction and isopropanol precipitation and diluted to 160ng/μl for microinjection.

*In vitro* fertilisation (IVF) was performed as described above (Sztein et al., 2000 (Sztein et al., 2000)) with minor modifications. Briefly, sperm were collected from the vasa deferentia and then capacitated for 1.5 hours in human tubal fluid (HTF) medium (Millipore). Sperm cells were then incubated with intact cumulus-oocyte complexes or zona-free oocytes for 5 hours. After removing excess sperm and cumulus lysate, presumptive fertilized eggs were transferred to equilibrated KSOM+AA medium (Millipore) for further culture and analysis.

### Immunohistochemistry (IHC)

IHC was carried out according to standard protocols. Briefly, ovaries from 4-8-week-old mice were fixed in 4% paraformaldehyde overnight at 4°C then washed for 3 x 10 minutes in PBS followed by transfer to 70% ethanol before being processed into paraffin wax blocks. 5μm sections were cut and mounted onto SuperfrostPlus slides (Thermo Fisher Scientific). After de-waxing and rehydration, slides were subject to antigen retrieval by pressure cooking at maximum pressure for 5 minutes in 0.01M sodium citrate buffer pH6.

To reduce non-specific staining from endogenous peroxidases, sections were incubated in 3% hydrogen peroxide (Sigma) solution for 15 minutes.

Prior to incubation with the primary antibody, incubation with normal serum and BSA was carried out to reduce non-specific binding. The brown signal was revealed after incubation with ImmPRESSTM HRP Anti Rabbit IgG (Peroxidase) Polymer detection kit followed by ImmPACTTM DAB Peroxidase (HRP) substrate (both from Vector laboratories).

The sections were counterstained with haematoxylin and imaged on a Zeiss AxioImager Z1.

### Immunofluorescence

Immunofluorescence (IF) experiments were performed as previously reported (Lin et al., 2014). Most oocytes/embryos were directly fixed in 4% paraformaldehyde for 15 minutes. To remove unbound histones some oocytes were incubated with pre-extraction buffer containing 0.5% Triton X-100 for 5 minutes, following Hajkova and Nashu’s protocol (Paull et al.), before being fixed in 4% paraformaldehyde. Fixed samples were permeabilised by 0.5% Triton X-100 and then blocked by 5% BSA before incubation with primary antibodies. The antibodies used were Hira (LSBio LS-C137477), Ubn1(LSBio LS-B11373), H3.3(Abnova H00003021-M01), Cabin1(Abcam ab3349), H3K9me2 (Abcam ab1220), H3K9me3 (Abcam ab8898). Secondary antibodies used were donkey anti rabbit IgG, 568nm (Thermo Fisher A10042), and donkey anti mouse IgG, 488nm (Thermo Fisher A21202). After washing and counterstaining with DAPI (Life Technologies D3571) samples were mounted using Vectashield® (Vector laboratories).

Images of stained oocytes/embryos were acquired by a spinning disk confocal (CSU-W1, Yokogawa) on an upright microscope frame (BX-63, Olympus) using a 30 or 60x silicon oil immersion objective (UPLSAPO 60XS2, Olympus) (Percharde et al., 2018). IF signal intensity was quantified using Fiji software.

### Proximity Ligation Assay

Proximity ligation assay was performed according to the instructions of the PLA Duolink® assay kit manual (Sigma). Briefly, wild type zygotes (2- and 4-hours post IVF) were fixed for 15 minutes in 4% paraformaldehyde followed by permeabilisation for 20 minutes in 0.5% TritonX in PBS. The samples were then blocked in PLA Duolink® blocking solution before being incubated with primary antibodies: either mouse anti Hira (LSBio LS-C137477) and rabbit anti Ubn1(LSBio LS-B11373) or mouse anti Hira and rabbit anti Cabin1 (Abcam ab3349). As a negative control the antibodies raised in rabbit were incubated with mouse anti hCG (Abcam ab9582) which should not be present in mouse tissues.

All wash steps were with Wash Buffer A. After washing, the samples were incubated with the PLA PLUS and MINUS probes, diluted as directed, for 60 minutes at 37°C. Following washing the samples were incubated for 30 minutes at 37°C in ligation reaction mixture followed by further washes before being incubated for 100 minutes at 37°C in the reaction mixture for rolling circle amplification. The samples were mounted on SuperfrostPlus® slides (Thermo Fisher Scientific) using Duolink® mounting medium plus DAPI. Images were acquired by a spinning disk confocal (CSU-W1, Yokogawa) on an upright microscope frame (BX-63, Olympus) using a 30 or 60x silicon oil immersion objective (UPLSAPO 60XS2, Olympus).

### Human one-pronucleus zygote collection

Approvals were obtained from an ethical committee (East of Scotland Research Ethics Service; REC 16/ES/0039) and the Human Fertilisation and Embryology Authority (HFEA), research licence (R0204), for the collection of human one-pronucleus (1PN) zygotes, in collaboration with the Edinburgh Fertility and Reproductive Endocrine Centre (EFREC). Patients were recruited randomly, and samples were collected from consented couples. Oocyte and sperm collection, fertilisation by IVF or ICSI, and embryo culture procedures, were performed following the routine operational protocols of EFREC and described in Sciorio et al., (Sciorio et al., 2018). Briefly, cumulus-oocyte-complexes (COC) were retrieved from follicular fluid, and then oocytes were isolated and washed. Oocytes were inseminated by IVF or ICSI as clinically indicated. Inseminated oocytes were cultured in G-IVF Plus medium (Vitrolife, Sweden) at 37 °C and 6% CO2 in atmospheric air. Single pronucleus zygotes were identified approximately 18 hrs post IVF or ICSI.

Embryo disposal, transfer, and witness documents followed the regulations of the HFEA clinic licence (0201). 1PN zygotes were individually labelled and transported using a portable incubator in G-MOPS Plus medium (Vitrolife, Sweden) at 37°C from the clinic laboratory to the research laboratory. Zygotes were then fixed in 4 % paraformaldehyde for 15 minutes at room temperature. IF procedure and confocal imaging were the same as described above. Antibody against H3K9me3 was used as a marker of human female chromatin and Ubn1, Hira, H3.3, and Cabin1 antibodies were also applied.

### Statistical analyses

For analyses of the percentage of pronuclear formation, the χ2 test was used. For analyses of IF intensity, two-tailed t-test with unequal variance was used. All error bars indicate s.d.

---

Figure S1. Interdependence of Hira complex molecules in the mouse oocytes.

(A) Experimental approaches to generate oocyte specific loss-of-function Hira complex models. Hira and Cabin1 mutant mice were generated by crossing Zp3-Cre conditional alleles with Hira flox/flox and Cabin1 flox/flox females respectively. Maternal depletion of Ubn1 was generated by microinjection of morpholino antisense oligos against Ubn1.

(B) Procedure of generation of Ubn1 knockdown oocytes and the assay to monitor male pronucleus formation. Ubn1 and control morpholino microinjected oocytes were matured in vitro to MII stage oocytes and then followed by fertilization. Zygotes were assessed after 5 hrs post fertilization for examination of male pronucleus formation.

(C) Ubn1 morpholino induced efficient knockdown. Immunofluorescence of Ubn1 of Ubn1 and control morpholino injected oocytes (left panel). Quantification of the result of immunofluorescence (right panel). Scale bar=40μm.

(D) Ubn1 loss-of-function oocytes revealed the decreased level of Hira and Cabin1. Immunofluorescence of Hira and Cabin1 in the Ubn1 knockdown oocytes (upper panel). Quantification of the results of immunofluorescence (lower panel). Scale bar=40μm.

(E) Cabin1 loss-of-function oocytes showed the decreased level of Hira complex and H3.3. Quantification result of the immunofluorescence of Cabin1, Ubn1, Hira, and H3.3 in the Cabin1 mutant oocytes (upper panel). Immunofluorescence (lower left panel) and quantification (lower right panel) result of the Cabin1 oocytes after Triton-X treatment showed that H3.3 failed to incorporate into Cabin1 mutant oocytes. Scale bar=40μm.

Figure S2. Validation of Hira overexpression in the Hira mutant oocytes. Immunofluorescence of Hira and H3.3 of the Hira mutant (ZH KO) oocytes after RNA injected (left panel). Quantification of immunofluorescence result showed that the level of Hira in the Hira mutant oocytes was comparable to heterozygous controls (ZH Het); right panel. Scale bar=80μm.

## Acknowledgement

This work was supported by MRC Centre Grant MR/N022556/1, the Wellcome Trust-University of Edinburgh Institutional Strategic Support Fund, Barbour Watson Trust, and grants from Deanery of Clinical Sciences, College of Medicine & Veterinary Medicine of University of Edinburgh to C.-J. L. C.-J. L. is a Royal Society of Edinburgh Personal Research Fellow funded by the Scottish Government.

We thank Anne Saunderson and Isobel Morton for recruitment of patients and management the clinical information. We also thank Martin Anger for the plasmid of GFP (McGuinness et al., 2009), Debasree Dutta for the plasmid of GFP-Hira (Majumder et al., 2015), and Peter Adams for the antibody of Ubn1.

We appreciate Profs Andrew Horne and Norah Spears for their constructive comments.

## Author contribution

R. S.: Mouse colony management and performed oocyte experiments.

S. P and A.K.: Performed human IVF treatment under the overall management of J.T.

R.A.A: HFEA Licence holder.

C.-J. L., R.S., and R.A.A. interpreted the data.

C.-J. L.: Conceived the project and designed the experiments performed all embryo micromanipulation and human IF work.

R.S., R.A.A, and C.-J. L. wrote the manuscript with input from all authors.

