## Supplementary material for "Histone H3.3 Hira chaperone complex contributes to zygote formation in mice and humans": Suppl Figures

### Slide 1
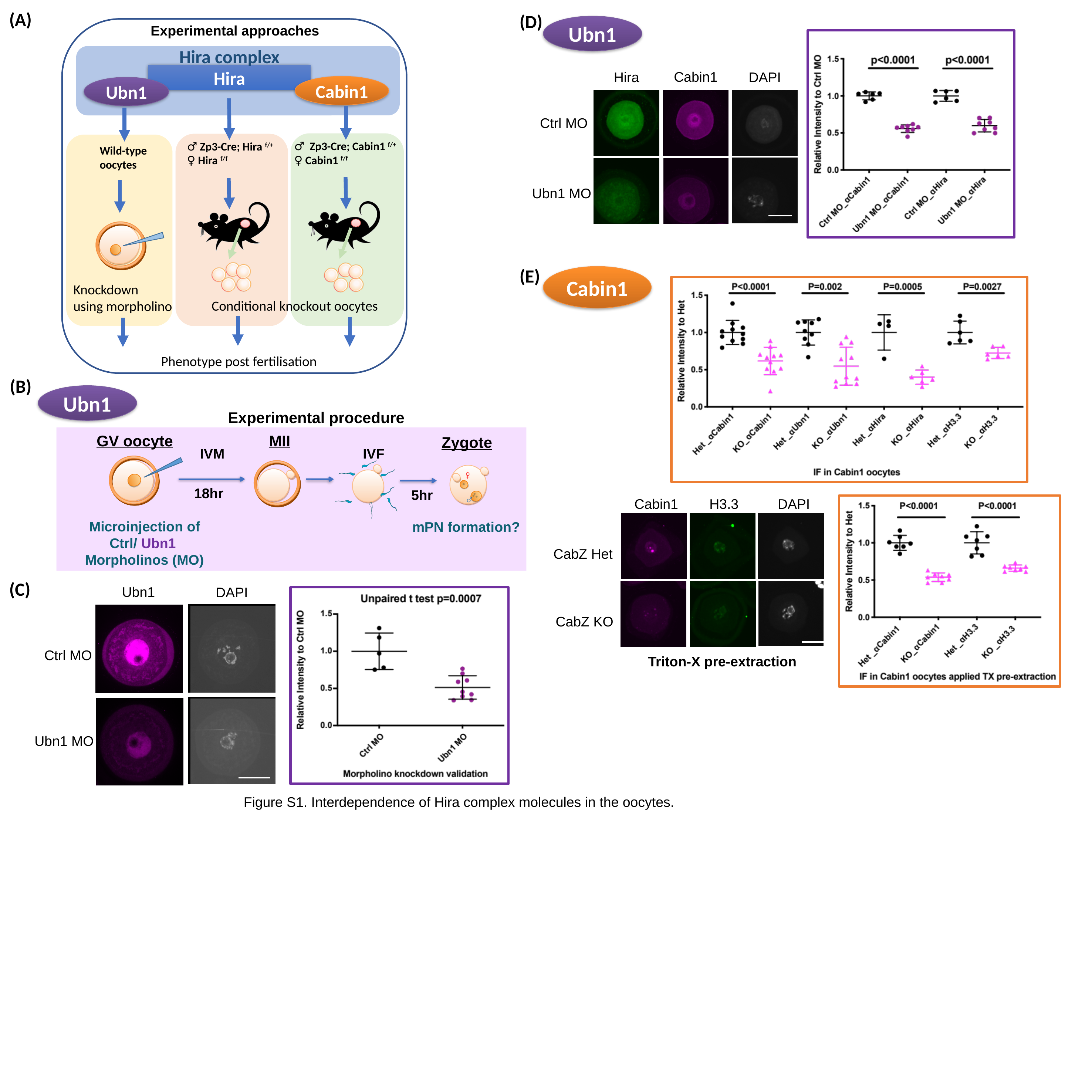

(A)
(D)
Cabin1
Hira
DAPI
Ctrl MO
Ubn1 MO
Ubn1
Experimental approaches
Hira complex
Hira
Cabin1
Ubn1
♂ Zp3-Cre; Cabin1 ^f/+^
♀ Cabin1 ^f/f^
♂ Zp3-Cre; Hira ^f/+^
♀ Hira ^f/f^
Wild-type
oocytes
Knockdown
using morpholino
Conditional knockout oocytes
Phenotype post fertilisation
(E)
Cabin1
DAPI
Cabin1
H3.3
CabZ Het
CabZ KO
Triton-X pre-extraction
(B)
Ubn1
Experimental procedure
GV oocyte
MII
Zygote
IVF
IVM
♀
18hr
5hr
♂
Microinjection of
Ctrl/ Ubn1
Morpholinos (MO)
mPN formation?
(C)
Ubn1
DAPI
Ctrl MO
Ubn1 MO
Figure S1. Interdependence of Hira complex molecules in the oocytes.

### Slide 2
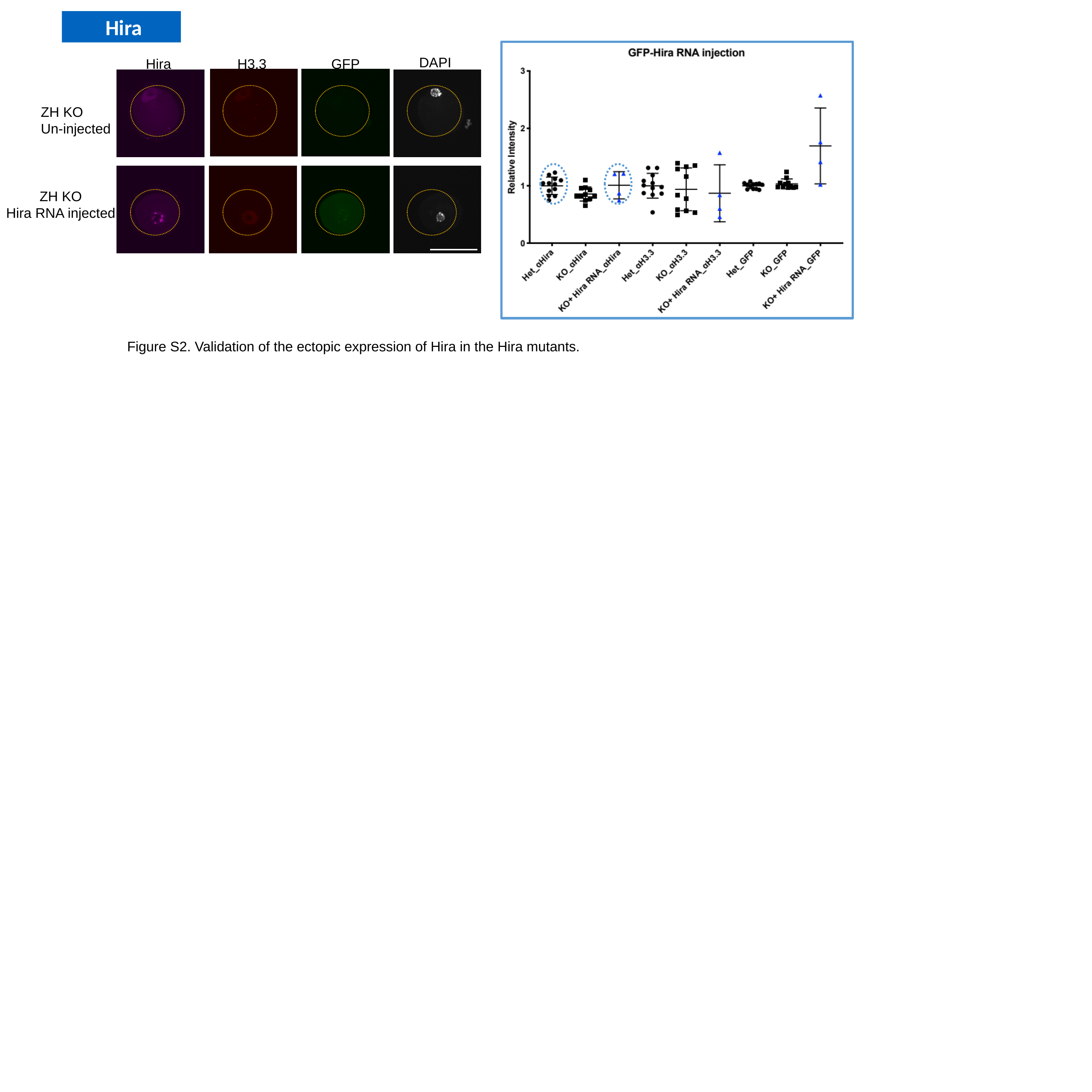

Hira
DAPI
Hira
H3.3
GFP
ZH KO
Un-injected
ZH KO
Hira RNA injected
Figure S2. Validation of the ectopic expression of Hira in the Hira mutants.
